# Role of U11/U12 minor spliceosome gene *ZCRB1* in Ciliogenesis and WNT Signaling

**DOI:** 10.1101/2024.08.09.607392

**Authors:** Geralle Powell-Rodgers, Mujeeb Ur Rehman Pirzada, Jahmiera Richee, Courtney F. Jungers, Sarah Colijn, Amber N. Stratman, Sergej Djuranovic

**Author notes:** these authors contributed equally to the work (Geralle Powell-Rodgers, Mujeeb Ur Rehman Pirzada, Jahmiera Richee).

## Abstract

Despite the fact that 0.5% of human introns are processed by the U11/U12 minor spliceosome, the latter influences gene expression across multiple cellular processes. The ZCRB1 protein is a recently described core component of the U12 mono-snRNP minor spliceosome, but its functional significance to minor splicing, gene regulation, and biological signaling cascades is poorly understood. Using CRISPR-Cas9 and siRNA targeted knockout and knockdown strategies, we show that human cell lines with a partial reduction in ZCRB1 expression exhibit significant dysregulation of the splicing and expression of U12-type genes, primarily due to dysregulation of U12 mono-snRNA. RNA-Seq and targeted analyses of minor intron-containing genes indicate a downregulation in the expression of genes involved in ciliogenesis, and consequentially an upregulation in WNT signaling. Additionally, *zcrb1* CRISPR-Cas12a knockdown in zebrafish embryos led to gross developmental and body axis abnormalities, disrupted ciliogenesis, and upregulated WNT signaling, complementing our human cell studies. This work highlights a conserved and essential biological role of the minor spliceosome in general, and the ZCRB1 protein specifically in cellular and developmental processes across species, shedding light on the multifaceted relationship between splicing regulation, ciliogenesis, and WNT signaling.

## INTRODUCTION

Pre-mRNA splicing—in which introns are excised and exons are fused together—is essential for gene expression, as the mRNA isoforms produced from this process are translated into proteins^1^. The spliceosome, a complex composed of several snRNPs (small nuclear ribonucleoprotein particles) and associated factors, catalyzes splicing through two consecutive transesterification reactions^2–5^. In most metazoans, two spliceosomes exist: the major (U2-dependent) and the minor (U12-dependent)^6,7^. The U2-dependent spliceosome removes over 99% of human introns, which are categorized as major (U2-dependent) introns, while the minor spliceosome removes less than one percent of a rarer class of human introns, minor (U12-dependent) introns^8^. Minor introns are distinguished from major introns by the absence of a polypyrimidine tract upstream of the 3′ splice site and the presence of a more tightly conserved 5′ splice site and branch point sequence (BPS)^9^. A major difference between the two spliceosomes is in their snRNP composition. The major spliceosome contains the snRNPs U1, U2, U4 and U6, while the minor spliceosome contains snRNPs U11, U12, U4atac, and U6atac. The only snRNP shared between both spliceosomes is the U5 snRNP^9–12^. Furthermore, fourteen unique proteins are specific to the minor spliceosome^13^. Over two decades ago, the first seven proteins associated with the U11/U12-disnRNP-U11/U12-20K (ZMAT5: Zinc Finger Matrin-Type 5), U11/U12-25K (SNRNP25: Small Nuclear Riboonucleoprotein U11/U12 Subunit 25), U11/U12-31K (ZCRB1: Zinc Finger CCHC-Type And RNA Binding Motif Containing 1), U11-35K (SNRNP35: Small Nuclear Ribonucleoprotein U11/U12 Subunit 35), U11-48K (SNRNP48: Small Nuclear Ribonucleoprotein U11/U12 Subunit 48), U11-59K (PDCD7: Programmed Cell Death 7, and U11/U12-65K (RNPC3: RNA Binding Region (RNP1, RRM) Containing 3),were identified by affinity purification and mass spectroscopy^12,14^. More recently, CENATAC (Centrosomal AT-AC Splicing Factor) was found to be the first protein specifically associated with U4atac/U6atac.U5 tri-snRNP^15^. It was also shown to have a minor spliceosome-specific binding partner, TXNL4B (Thioredoxin Like 4B)^15^. Purification and structural characterization of the human minor B^act^ complex by cryo-electron microscopy (cryo-EM) has identified additional minor spliceosome-specific proteins that contribute its stability and function -SCNM1 (Sodium Channel Modifier 1) and CRIPT (CXXC Repeat Containing Interactor of PDZ3 Domain)^16^ The study also confirmed the presence of RBM48 (RNA Binding Motif Protein 48) and ARMC7(Armadillo Repeat Containing 7), interacting partners shown to be required for minor splicing in maize^16,17^. An investigation in the post-spliceosomal complex, identified RBM41, a close paralog of U11/U12-65K^18^. Moreover, ZRSR2 (Zinc Finger CCCH-Type, RNA Binding Motif, and Serine/Arginine Rich 2), which has been shown to associate with both major and minor spliceosomes, has been shown to predominantly affect the splicing of minor introns^19^.

Although U12-dependent introns consist of a small percentage of human introns, the genes that contain them are enriched in several important information-processing events. Information processing events are the biological processes that involve the regulation and transmission of genetic information, such as DNA replication and repair, RNA transcription and translation, splicing, cytoskeletal organization, and various signaling pathways^6,20^. The most highly enriched functional process characterized for human MIGs (minor intron-containing genes) is cell-cycle regulation, with most MIGs identified as essential for cell division^21^. As such, multiple minor spliceosome-specific genetic deficiency models have been studied to assess the role of minor introns in gene expression and eukaryotic development. *RNPC3* (RNA binding region (RNP1, RRM) containing 3), which encodes the minor spliceosome-specific U11/U12-65K protein, has been extensively studied in eukaryotic models and is essential for the formation and stability of the intron recognition competent U12/U11 di-snRNP through its direct binding of the U12 snRNA and the U11-59K protein^22^. Loss of the zebrafish ortholog of *RNPC3*, *rnpc3*, leads to altered global gene expression and minor intron retention, with enrichment in differential expression and retained introns in genes involved in mRNA processing, splicing, transcription, and nuclear export^23^. Additionally, zebrafish mutants with loss of *rnpc3* showed gross morphological abnormalities during organogenesis, particularly in the development of the eyes and endodermal organs. Loss of *rnpc3* was lethal, with fish dying between 7 to 10 days post fertilization^23^. Studies in Arabidopsis show the U11/12-65K and U11/12-31K homologs are indispensable for plant growth, development, and minor intron splicing, with global analysis of u*11/12-65K* mutants showing mis-splicing of roughly 85% of minor-intron containing genes^24,25^. These findings underscore the critical importance of minor spliceosome-associated proteins and the minor spliceosome in early development.

Moreover, loss of function mutations in minor spliceosome-associated factors have implications for human disease. There are several congenital human diseases that have been shown to occur, explicitly as a result of the dysregulation of one or more of the integral U12-dependent snRNP components: *RNU4ATAC*-RFMN (Roifman syndrome), MOPD1/TALS (microcephalic osteodysplastic primordial dwarfism type I/Taybi–Linder syndrome), LWS (Lowry Wood syndrome), and JBTS (Joubert syndrome); *RNU12*-EOCA (early onset cerebellar ataxia) and CDAGS syndrome; *RNPC3*-IGHD (isolated growth hormone deficiency); and CENATAC-MVA (Mosaic Variegated Aneuploidy).^15,26–28^. Other disorders resulting from mutations in U12 spliceosome-associated proteins include Orofaciodigital syndrome (OFD) caused by mutations in *SCNM1*, a Rothmund-Thomson-like syndrome caused by mutations in *CRIPT*^29–31^. Somatic mutations in ZRSR2 have been shown to be causative in Myelodysplastic Syndrome (MDS) and associated with an increased risk of progression to Acute Myeloid Leukemia (AML)^19,32^ Additionally, the regulated expression of minor spliceosomal components and factors has also been implicated in the etiology and progression of multiple cancers^33,34^. However, these lists are not exhaustive, and ongoing discoveries about minor spliceosome factors are likely to reveal additional associated conditions.

Despite the importance of the minor spliceosome in cellular homeostasis and human disease, the molecular and functional mechanisms by which key factors regulate cellular homeostasis remains to be elucidated. One of these is the minor spliceosome-specific protein ZCRB1 (zinc finger CCHC-type and RNA binding motif containing 1). Although ZCRB1 associates with both the U12 mono-snRNP and U11/U12 di-snRNP (responsible for intron recognition), the mechanism by which it affects minor splicing is unknown. Transient knockdown of ZCRB1 results in the retention of minor introns, yet the functional significance of ZCRB1-mediated splicing, and the interplay between this role and other biological processes, also remains unknown^35^. Finally, functional genomic analysis of human cancer cell lines identified the essentiality of ZCRB1, necessitating a need to study its fundamental role in cellular homeostasis and survival ^36,37^.

In this study, using CRISPR based gene knockdown models in human cells and zebrafish - combined with functional genetics, biochemistry, and transcriptomic analysis - we show that loss of ZCRB1 in both humans and zebrafish leads to altered expression of minor intron containing genes essential for primary cilia formation and maintenance. Primary cilia, non-motile organelles present on the surface of most vertebrate cells, are comprised of a microtubule-based axoneme anchored by a basal body (mother centriole) and are crucial for cellular signaling and cell polarity during development ^38^. As such, in human cells and our zebrafish model with ZCRB1 loss, we see dysregulated primary cilia and upregulation of WNT-signaling. The dysregulation of WNT signaling is coupled with body axis defects and failed gastrulation in zebrafish—effects that can be rescued by the re-expression of wild type (WT) human *ZCRB1* or by introduction of a WNT inhibitor. In contrast, while loss of cilia is rescued by re-expression of human *ZCRB1* in *zcrb1* mutant zebrafish, using a WNT inhibitor does not rescue ciliogenesis. Our results provide a new role and model for the minor spliceosome and its associated gene *ZCRB1* in the maintenance of primary cilia gene expression, the control of WNT signaling, and its essential role in vertebrate development.

## RESULTS

### *ZCRB1* is an essential gene and partial loss of *ZCRB1* stimulates WNT signaling in Hek293 cells

To determine the effects of ZCRB1 loss-of-function on cellular homeostasis and the cellular transcriptome, we employed precise CRISPR/Cas-9 genome editing using two independent rounds of sgRNA transfection in Hek 293 Flp-In T-Rex cells (Figure 1A) to introduce a GFP donor fluorophore and targeted genomic mutations. The GFP donor persists extra-chromosomally, functioning as an episomal marker of gene editing. Following Fluorescence Activated Cell Sorting (FACS) for GFP-positive cells, we expanded 80 single cell colonies, of which 48 edited lines survived. Whole Genome Sequencing (WGS) of eight of these lines, selected at random, identified the creation of a *ZCRB1* heterozygous line harboring a 1 bp insertion in the first exon of *ZCRB1* (base42324038; Chr12:42324021-4232041). This deletion led to a +1 frameshift and early termination of translation at amino acid residue p.Thr22fs. (Fig 1a., Table 1., Supplementary Fig. 1). As all of the selected clones were heterozygous for *ZCRB1*, we selected three—33, 63, PL3C7—to cut with a second sgRNA in an independent region, and obtained complex *ZCRB1* homozygous deletion cells (Table 1). The second sgRNA was designed to target the fourth coding exon (Chr12:42317406-Chr12:42317426, negative strand, Fig. 1a). Despite two attempts to generate *ZCRB1* homozygous mutant cell lines, we were only able to produce heterozygous mutants (Fig. 1b-c, Table 1), suggesting that homozygous deletion of *ZCRB1* is lethal. However, in targeting these heterozygous mutants with a second sgRNA, we created an independent line harboring a 14 bp deletion (base 42317399) that induced a frameshift (p.Ile87fs (Figure 1a, Table 1, Supplemental Fig. 1). This deletion resides on the same allele as the 1 bp deletion, again leading to only monoallelic loss of *ZCRB1*. This 14 bp edit was only found in Clone 54 (Table 1, Supplemental Fig. 1). We used CassOFFinder software, in conjunction with WGS, to detect off target events due to genomic editing by the two sgRNAs (Supplemental Table 1a)^39^. Out of more than 1000 events for each sgRNA, we found only the predicted benign intronic variant in DAB1 (Disabled-1; c.-137+177702T>C) from genomic editing with gRNA 1 (Supplemental Table 1b), supporting minimal off-target effects from the selected gRNAs. We tested 16 cell lines derived from the individual *ZCRB1* clones for expression of ZCRB1 protein (14 edited and 2 non-edited clones; Extended Data Fig. 1). The 14 edited and confirmed heterozygote clones reduced ZCRB1 protein levels while the two CRISPR/Cas9 targeted but non-edited clones (B7 and C7pl1) had ZCRB1 protein levels similar to parental HEK293 cells (Fig. 1b, Extended Data Fig 1). We selected four of the edited clones and confirmed a partial loss of *ZCRB1* transcript levels, as measured by qRT-PCR (Fig. 1c). Each CRISPR/Cas-9 selected clone had a 30-50% reduction in steady-state *ZCRB1* mRNA levels, with the greatest reduction in Clone 54 (65%) compared to WT (Fig. 1c). Despite multiple attempts to create complete *ZCRB1* knockout cells, we only recovered clones carrying monoallelic losses of *ZCRB1*, suggesting its essential role in cellular fitness and survival.

**Figure 1:**
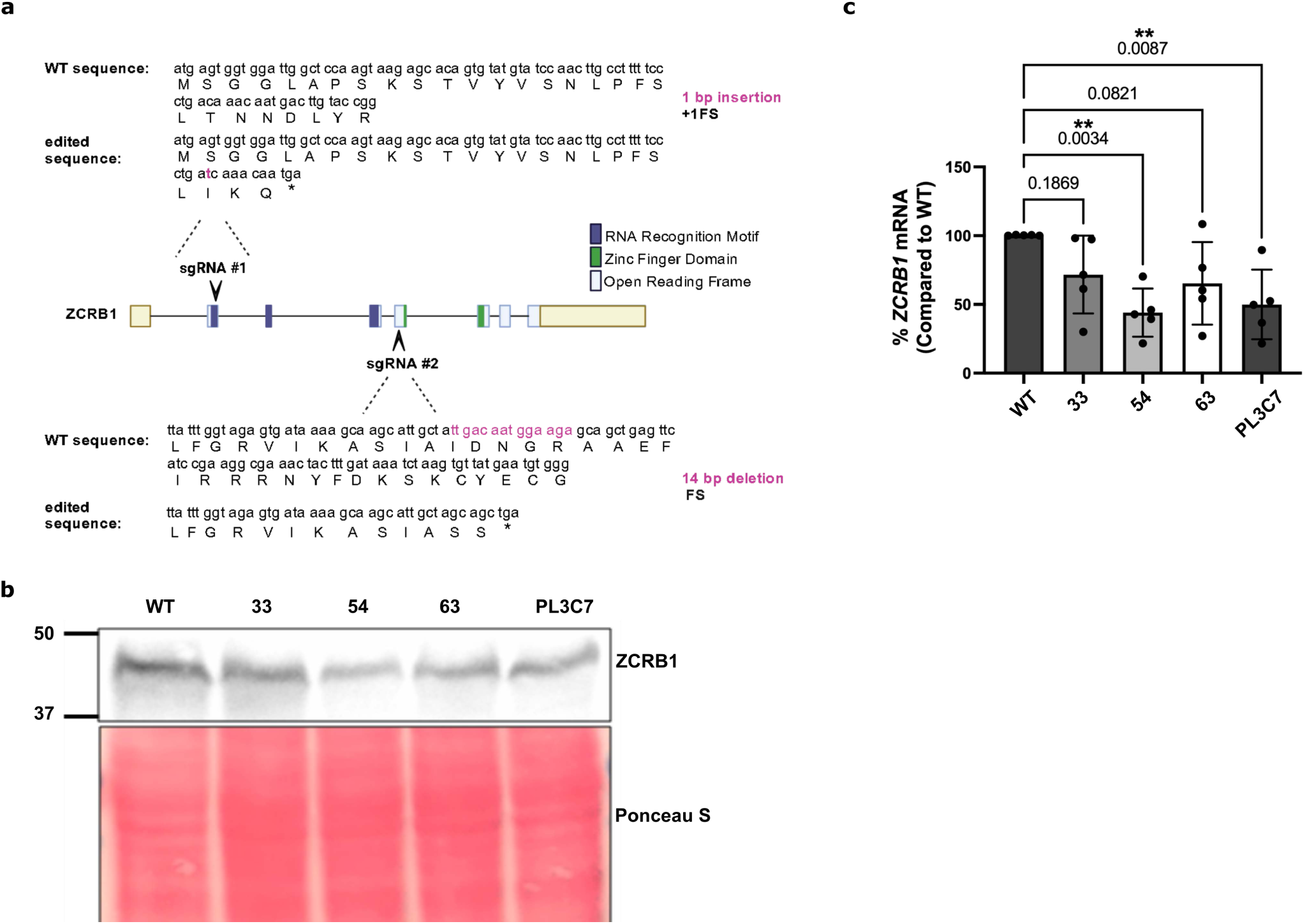
*ZCRB1* is an essential gene in Hek293 cells a. Schematic representation of CRISPR/Cas9 editing of the ZCRB1 gene in Hek293 Flp-In cells. The domain structure of ZCRB1 is depicted, with the locations of two distinct sgRNAs indicated by black arrows. The genomic and translated amino acid sequences are shown for both the unedited wild-type (WT) and edited versions. The specific genomic edits and their corresponding outcomes are highlighted. b. Western blot analysis of Hek293 Flp-In cell lines with heterozygous loss of ZCRB1. ZCRB1 expression is shown for WT Hek293 Flp-In cells and 33, 63, 54, 63, and PL3C7 clones, respectively. Molecular weight markers (kDA) are shown. Equal protein was used for analysis and was normalized using Ponceau S staining as shown. c. ZCRB1 mRNA levels of ZCRB1 heterozygous knockout clones: 33, 63, 54, and PL3C7, respectively, in comparison to WT Hek293 Flp-In cell levels as quantified by qRT-PCR. Relative mRNA levels for each clone are shown as a percentage with respect to WT. Error bars represent mean ± s.d. values (n=3). Numbers above brackets represent the p-value as measured by an ordinary one-way ANOVA with Dunnett’s multiple comparisons test (omnibus p-value=**0.0085).

**Table 1:**
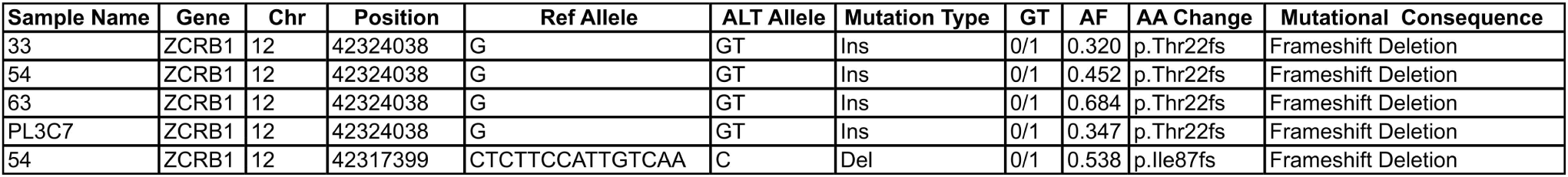
Gene dosage and functional consequences of CRISPR/Cas9-mediated editing of ZCRB1.

To determine the role of ZCRB1 in the regulation of gene expression, we performed RNA sequencing on the HEK293 parental cell line and four GFP+ selected heterozygous mutants, followed by differential gene expression analysis. After controlling heteroscedasticity in all samples, we saw a 2.6-fold reduction of *ZCRB1* transcript levels in the *ZCRB1* mutants compared to the WT samples (Supplemental Table 2a). Further analysis revealed significant dysregulation of 4096 protein-coding genes (2185 were upregulated and 1911 were downregulated) between WT samples and the four mutant clones. There were also 377 (227 upregulated, 150 downregulated) significant differentially expressed noncoding RNAs, which were primarily long non-coding RNAs (346/377; Figure 2a, Supplemental Table 2a). Gene set enrichment analysis performed by GAGE (Generally Applicable Gene-set Enrichment) identified high confidence (FDR ≤ 0.05) downregulated pathways involved in RNA metabolism and metabolic processes in *ZCRB1* mutant cells, while biological processes involving cell signaling and tissue and organ morphogenesis were upregulated (Extended Data Fig. 2, Supplemental Table 2b)^40^. Several snRNA genes that comprise the major and minor spliceosome, such as U1 and U12 snRNP, were downregulated. Moreover, 3 of the 8 U11/U12 specific di-snRNP-associated proteins were altered in their expression in addition to *ZCRB1*: *SNRNP25, SNRNP48* and *ZRSR2*. There was significant enrichment in downregulated genes belonging to the snRNP core component Sm gene family and those belonging to the SF3b complex, which is responsible for branch point sequence binding in both spliceosomes (Fig. 2b, Supplemental Table 2c)^41^.

**Figure 2:**
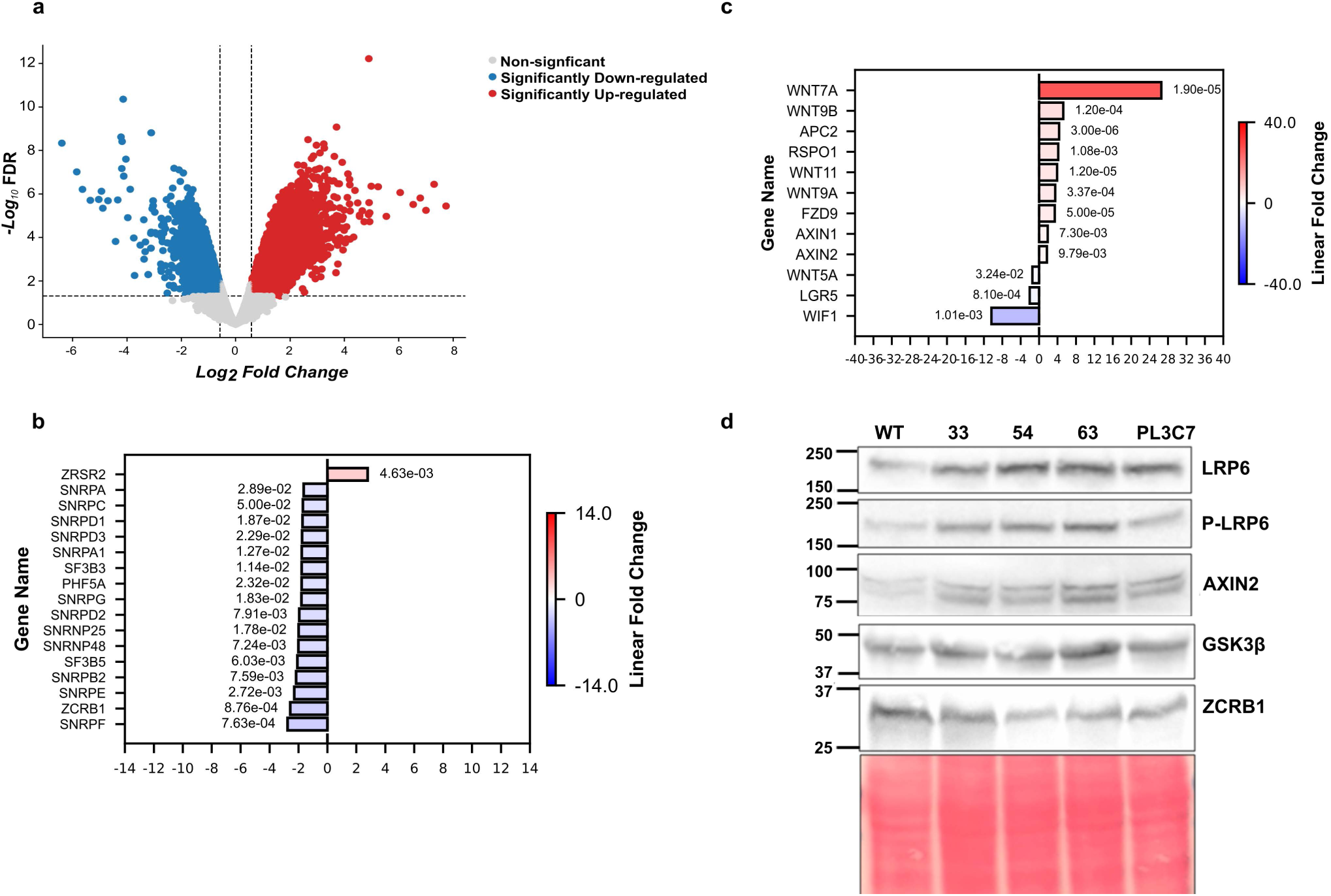
Partial loss of ZCRB1 leads to downregulation of splicing factors and upregulation of Wnt signaling a. Volcano plot of differentially expressed (DE) genes identified by RNA-seq analysis between heterozygous ZCRB1 knockout Hek293 Flp-In cells and WT cells. The red and blue dots denote significantly upregulated and downregulated genes, respectively, while the grey dots denote genes with no measure significant change. The horizontal dashed lines represent the cut off for significant fold change (≥ absolute log2 fold change of .58), while the horizontal line signifies the FDR significance cut-off (≤ 0.05). b. Barplot of DE genes belonging to or primarily associated to both the major and minor Human spliceosomes (FDR ≤ 0.05 and absolute linear fold change ≥ .1.5) as identified by RNA-seq analysis. Red and blue bars, upregulated and downregulated linear foldchanges, respectively. The color intensity denotes the degree of expression change. The FDR values for each expression level are displayed next to their respective bars. c. Barplot of DE genes present in the canonical and non-canonical Human Wnt pathway (FDR ≤ 0.05 and absolute linear fold change ≥ .1.5) as identified by RNA-seq analysis. Red and blue bars, upregulated and downregulated linear foldchanges, respectively. The color intensity denotes the degree of expression change. The FDR values for each expression level are displayed next to their respective bars. d. Western blot analysis of Wnt signaling factors in ZCRB1 heterozygous knockout cells. Protein for each labeled protein is shown for WT HEK293 flp-In cells and 33, 63, 54, 63, and PL3C7 clones, respectively. Molecular weight markers (kDa) are shown. Equal protein was used for analysis and was normalized using Ponceau S staining as shown.

Gene ontology (GO) analysis also revealed terms associated with developmental processes with 38 genes in the canonical and non-canonical WNT/Ca2+ and planar cell polarity (PCP) pathways regulated (Supplemental Table 2b,d) in response to *ZCRB1* partial loss-of-function. We found an increase in the expression of genes in the canonical WNT pathway: most strikingly *Wnt7a* was upregulated 26.6-fold, followed by *Wnt9B* and *Wnt11*, with 5.3 and 4.9-fold upregulation in expression, respectively (Fig. 2c). *Axin1* and *Axin2*, regulators of beta-catenin and WNT signaling, were also upregulated by 1.9 and 1.7-fold, respectively. We also observed a downregulation of inhibitors of canonical WNT signaling such as *Wif1* (10.3-fold reduction in *ZCRB1* mutant cells). These data suggested an upregulation of WNT signaling following partial loss of *ZCRB1* (Fig. 2b,c). Further analyses of the WNT signaling pathway at the protein level indicated upregulation of phosphorylated LRP6—a downstream WNT target gene—in the *ZCRB1* mutant cells compared to WT cells (Fig. 2d). Similarly, we saw increased protein levels of Axin2 in the mutant cells (Fig. 2d), consistent with our RNA-seq analyses and demonstrating that core components necessary for activation of canonical WNT signaling are upregulated following ZCRB1 loss-of-function.

### ZCRB1 loss downregulates cilia genes via improper splicing of minor introns

To further investigate how heterozygous loss of *ZCRB1* stimulates WNT signaling, we sought to determine whether changes in gene expression were specific to genes that harbored U12-dependent introns. The number of significant and differentially regulated genes that contained at least one U12-dependent intron was 164 of a total of 4582 genes that all had U2-dependent introns in our RNA-seq dataset. As such, genes with minor introns were affected overall at a slightly higher percentage: 28.3% of total minor intron-containing genes (164/579) were significantly affected by *ZCRB1* loss-of-function compared to 26.9% of genes with U2-dependent introns (4582/17,037). Moreover, we saw a significant enrichment in downregulated MIGs in comparison to downregulated non-MIGs (Fisher’s exact test, p-value = 6.6×10^−13^). In total, there were 121 downregulated and 43 upregulated minor intron genes with at least 1.5-fold change in the mRNA levels compared to 2082 downregulated and 2499 upregulated non-MIG genes, respectively (Fig. 3a, Supplemental figure 2a). Gene set enrichment analysis of these 164 differentially expressed minor intron genes found a statistically significant enrichment of processes involved in the assembly and regulation of primary cilia (Fig. 3b, Supplemental Table 3a).

**Figure 3:**
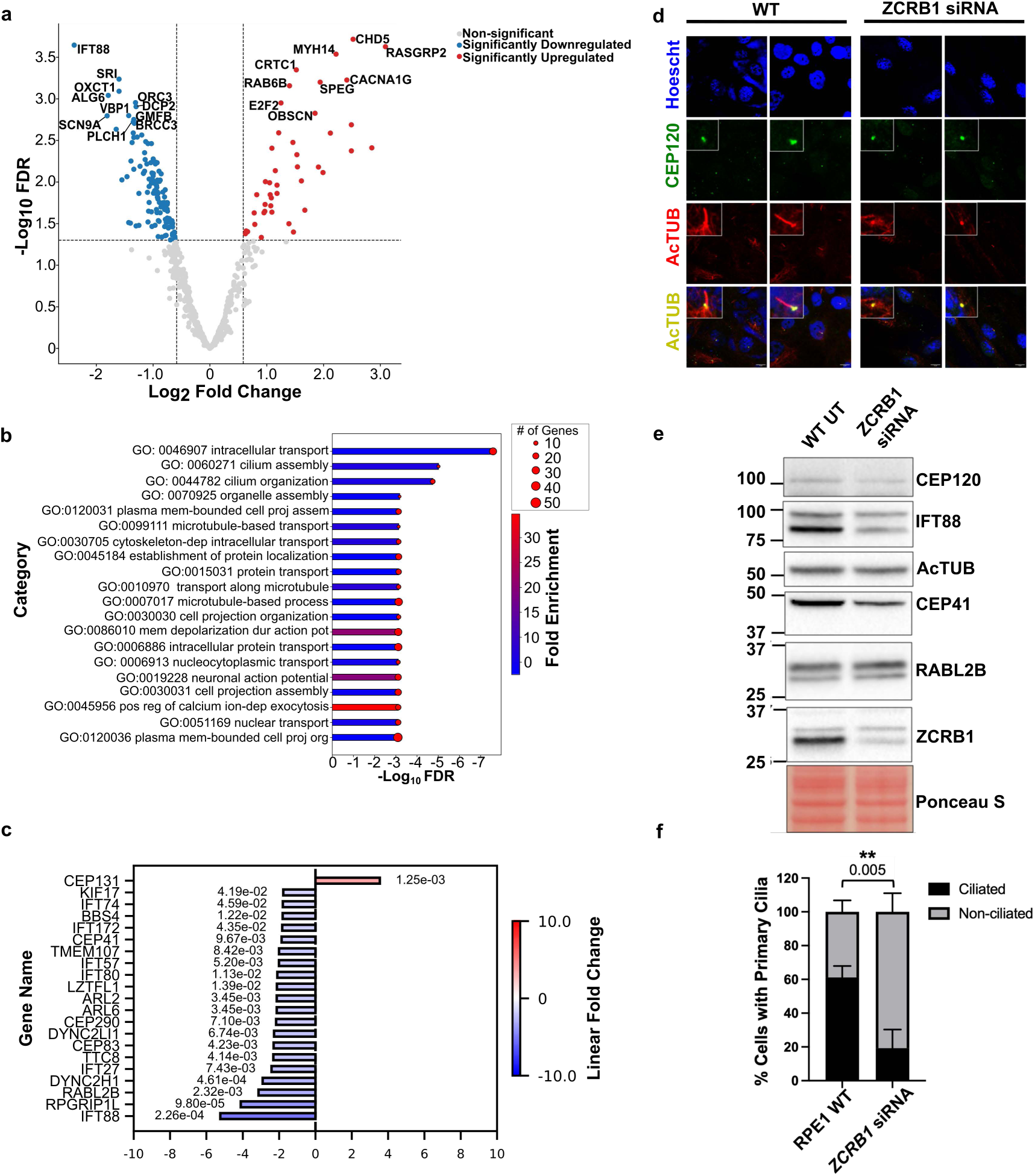
ZCRB1 regulates ciliogenesis genes via splicing of minor introns a. Volcano plot of DE MIG identified by RNA-seq analysis between heterozygous ZCRB1 knockout HEK Flp-in cells and WT cells. The red and blue dots denote significantly upregulated and downregulated genes, respectively, while the grey dots denote genes with no measure significant change. The horizontal dashed lines represent the cut off for significant fold change (≥ absolute log2 fold change of .58), while the horizontal line signifies the FDR significance cut-off (≤ 0.05). b. Barplot representative of the top FDR significant enriched biological processes (BP) identified involving DE MIGs in heterozygous *ZCRB1* knockout cell lines versus WT. The length of bars is proportional to the FDR (adjusted p value) and the shading dictates the degree of linear fold change shown by the color bar. The diameter of the red dots on the end of the bars represents the number of genes present in each identified enriched pathway. c. Barplot representative of top DE primary cilia and centrosome associated MIGs. (FDR ≤ 0.05 and absolute linear fold change ≥ .1.5) as identified by RNA-seq analysis. Red and blue bars, upregulated and downregulated linear foldchanges, respectively. The color intensity denotes the degree of expression change. The FDR values for each expression level are displayed next to their respective bars. d. Immunofluorescence for centrioles (CEP-120) and PC (acetylated tubulin) e. Western blot analysis of primary cilia associated proteins following siRNA-mediated ZCRB1 knockdown in RPE-1 cells. Protein expression of target genes is shown for WT and ZCRB1 siRNA targeted genes, respectively. f. Barplot showing quantification of primary cilia abundance. Statistics were calculated using an unpaired t test.

Furthermore, we observed significant differential expression of 116 genes involved in centrosome and primary ciliogenesis and function, of which 77 of the genes are downregulated. These genes included intraflagellar transport (IFT) proteins, which are necessary for cilia assembly, length, and maintenance as their central role involves the bi-directional movement of protein cargo along the axoneme^13^. Members of both the A and B IFT complexes, which control anterograde and retrograde transport in concert with kinesin-2 and dynein-2, respectively, were affected^31^. Of the 16 associated IFT-B complex proteins, 6 were downregulated in our differential expression analysis (*IFT88, IFT172, IFT74, IFT80, IFT57, IFT27*). *IFT88,* which is a core component of the IFT-B complex, is decreased 5.6-fold compared to levels in WT cells (Fig. 3c, Supplemental Table 3a)^42–44^.

In addition to the IFT machinery, we observed dysregulation of Bardet-Biedl syndrome complex members (BBSome), including *TTC8, ARL6, LZTFL1*, and *BBS4*. The BBSome interacts with IFT complexes to transport signaling molecules to the ciliary membrane^45^. Among the 116 centrosome and primary cilia-related genes analyzed, we identified downregulation of ten MKS (Meckel-Gruber syndrome) genes (*TMEM218, RPGRIP1L, TMEM17, TCTN1, TMEM231, TMEM107, TCN3, TMEM67, B9D1, TCTN2*) and genes associated with NPHP (nephronophthisis) (*NPHP3, CEP290, INVS, IQCB1*) (Supplemental Table 3a)^46^. These genes are crucial for maintaining the linkage between the basal body and the ciliary membrane, necessary for axoneme extension, intraflagellar transport, and the regulation of protein movement between the cilium and cytoplasm^46,47^ We also performed differential splicing analysis using RMATs^48^. After filtering, we found numerous statistically significant (FDR <0.05) alternative splicing events. Skipped exon events were the most abundant at 1907 events, followed by 644 mutually exclusive exon events, 382 retained intron events, 310 alternative 5’ start site events, and 258 alternative 3’ start site events (Supplemental Table 3b, Extended Fig. 3). Intron retention analysis showed a small but significant enrichment in minor intron genes (Fischer exact, p-value = 0.03) (Supplemental Table 3b, Extended Fig.3). The 29 retained intron events were associated with 20 unique minor intron genes. Of these 20 genes, minor introns were specifically retained for multiple primary cilia associated genes, including *CCDC28B*, a conserved protein involved in the maintenance of ciliary length (Supplemental Fig. 2b) ^49^. To validate these findings, we analyzed the effects of *ZCRB1* partial loss-of-function on primary cilia (PC) formation in retinal pigment epithelial cells-1 (RPE-1) that form PC more frequently and prominently compared to HEK293 cells^50^. Following *ZCRB1* siRNA treatment for 100 hrs, we observed a decrease in PC and protein levels containing minor introns critical for PC formation, including IFT88, RABL2B, and CEP41 (Fig. 3 d,e). Immunofluorescence labeling of centrioles (CEP-120) and PC (acetylated tubulin) confirm a significant decrease (80%) in PC formation in *ZCRB1* siRNA treated cells (Fig. 3 d,e,f). These results revealed a role for *ZCRB1* in positively regulating ciliogenesis via unerring splicing of pro-cilia related MIGs.

### ZCRB1 is essential in zebrafish development and controls Wnt signaling

In parallel to our investigations on the biological significance of ZCRB1 in human cells, we employed CRISPR/Cas12a genome editing in zebrafish embryos to specifically study the role of *zcrb1* in regulating gene expression during developmental tissue patterning^51^. The human and zebrafish Zcrb1 proteins share substantial sequence homology (72%) with an even higher degree of similarity in the RNA recognition motif (RRM) and the zinc finger (ZF)-CCHC type domains, 76% and 94% respectively (Fig. 4a). Transient mosaic Crispant (F0) embryos generated by targeting *zcrb1* in the fourth coding exon, led to a 1.4-fold reduction in *zcrb1* transcript expression compared to Cas12a-only injected siblings (Fig. 4b, Supplemental Table 4a). Phenotypic analysis of 28-hour post-fertilization (hpf) embryos showed 55% of the *zcrb1* Crispant embryos failed to gastrulate, 20% of injected fish were wild-type presenting, and the final 25% presented with disrupted dorsal-ventral body axis patterning (Fig. 4c). The gastrulation and body axis phenotypes could be rescued by reintroduction of human WT *ZCRB1* mRNA co-injected into the embryos at the 1-cell stage with the *zcrb1* CRISPR gRNA, suggesting the phenotypes noted are specific to *zcrb1* gene editing in the zebrafish (Fig. 4c). To study the effects of partial loss of Zcrb1 function on gene expression in the zebrafish, we performed RNA-seq and differential gene expression analysis on Crispant injected (F0) embryos compared to Cas12a only injected WT control siblings. From this analysis, we observed global dysregulation in gene expression, with significant downregulation of 3408 genes and significant upregulation of 6199 genes (Fig 4d, Supplemental Table 4a). Gene set enrichment analysis identified gene programs upregulated in biological processes including cellular components biogenesis, RNA processing, and cellular and organismal development (Fig 4e, Supplemental Table 4b). Similar to observations in human cells heterozygous for *ZCRB1* expression, we note significant differential expression of several members of the canonical and non-canonical Wnt signaling pathways. Of the 120 differentially expressed Wnt signaling pathway genes, 16 were downregulated, while 104 were upregulated (Supplemental Table 4c). In particular, Wnt pathway inhibitors, including *wnt5a* and members of the dickkopf (*dkk*) family, were downregulated in *zcrb1* Crispant embryos, while positive regulators of the Wnt signaling pathway, such as *wntt11f2* (paralogue of human *Wnt11*), were upregulated (Fig. 4f, Supplemental Table 4c), consistent with the data seen in edited HEK293 human cell lines. Given the high conservation between human and zebrafish ZCRB1, the developmental effects observed in zebrafish attributed to partial *zcrb1* loss, our findings across both organisms highlight its role in Wnt signaling modulation.

**Figure 4:**
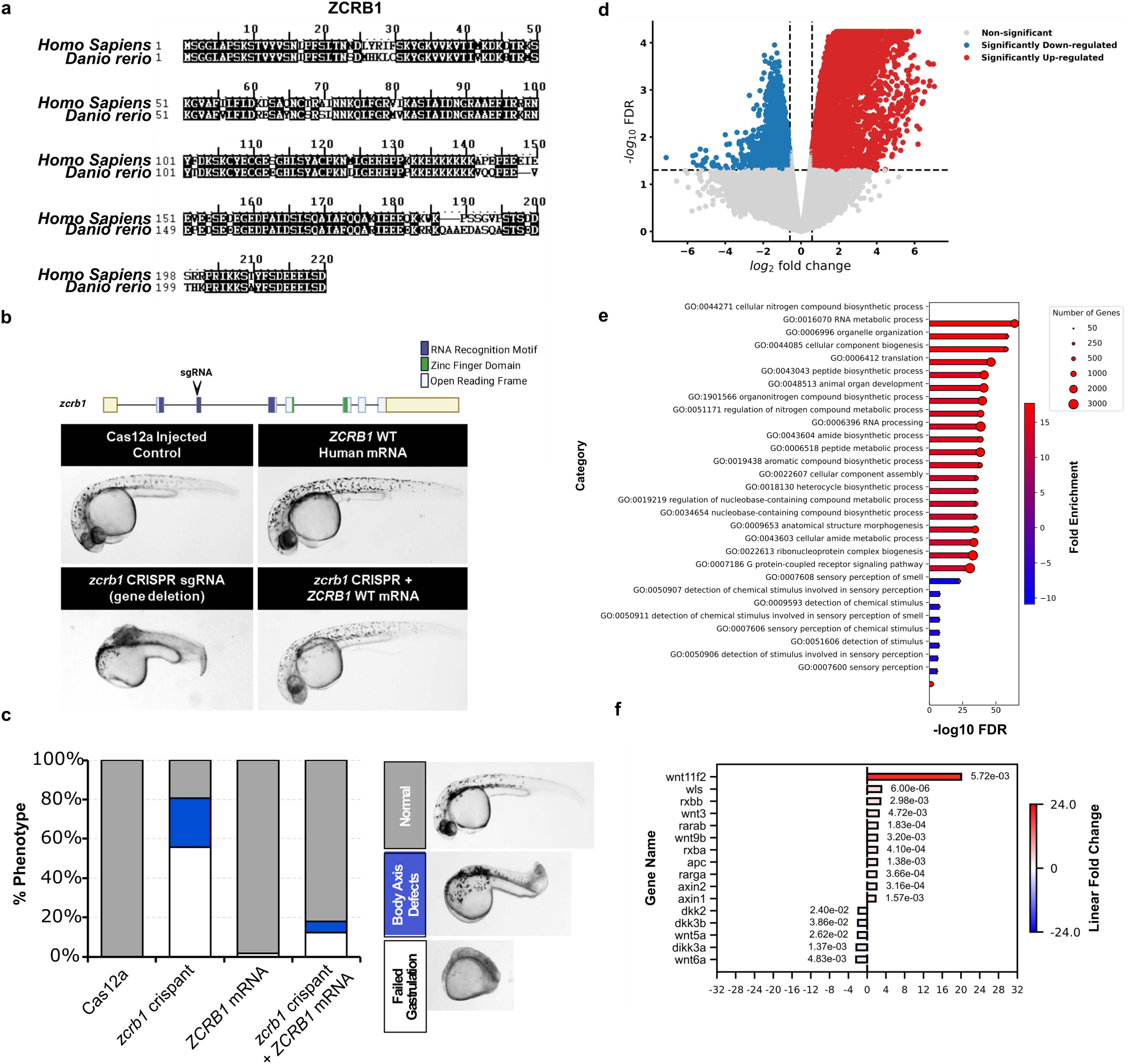
zcrb1 is essential for Wnt signaling and development in zebrafish a. Alignment of ZCRB1 amino acid sequence in human and zebrafish, respectively. Shaded regions represent exact homology. b. Phenotype analysis of CRISPR/Cas12a mediated editing of zcrb1 in zebrafish embryos. Schematic is representative of zcrb1 locus and location of CRISPR/Cas12a editing. c. Stacked barplot representing the percentage of phenotypes show in Cas12a and *zcrb1* crispant embryos, WT embryos injected with *ZCRB1* mRNA, and *zcrb1* crispant embryos injected with *ZCRB1* mRNA. d. Volcano plot representative of DE genes identified by RNA-seq between *zcrb1* crispant embryonic cells and WT. The red and blue dots denote significantly upregulated and downregulated genes, respectively, while the grey dots denote genes with no measure significant change. The horizontal dashed lines represent the cut off for significant fold change (≥ absolute log2 fold change of .58), while the horizontal line signifies the FDR significance cut-off (≤ 0.05). e. Barplot representative of the top FDR significant enriched biological processes (BP) identified in crispant zcrb1 embryonic cells vs WT. The length of bars is proportional to the FDR (adjusted p value) and the shading dictates the degree of linear fold change shown by the color bar. The diameter of the red dots on the end of the bars represents the number of genes present in each identified enriched pathway. f. Barplot representative of top DE Wnt signaling pathway genes (FDR ≤ 0.05 and absolute linear fold change ≥ 1.5) as identified by RNA-seq analysis. Red and blue bars, upregulated and downregulated linear foldchanges, respectively. The color intensity denotes the degree of expression change. The FDR values for each expression level are displayed next to their respective bars.

### Impaired cilia formation and upregulation of WNT signaling components is conserved in zebrafish following *zcrb1* loss of function

Our results in human RPE1 cells indicated impaired splicing and loss of primary cilia components as a primary consequence of decreased *ZCRB1* function (Fig. 3 d-f). To understand if loss of primary cilia formation was conserved in the zebrafish, we injected the gRNA against *zcrb1* into *Tg(actb2:Mmu.Arl13b-GFP)^hsc5Tg^*transgenic zebrafish, where all cilia are labeled with GFP for live imaging, to generate F0 Crispants^52^. Four conditions were assessed, Cas12a only injected; *zcrb1* Crispants; *ZCRB1* WT human mRNA injected; and *zcrb1* Crispants plus *ZCRB1* WT human mRNA injected together. Images acquired at 32 hpf across all four conditions showed cilia numbers were decreased in the *zcrb1* Crispant injected embryos and rescued nearly back to control levels following co-injection of *ZCRB1* WT human mRNA with the *zcrb1* Crispant gRNA (Fig. 5 a,b). Together these data suggested that impaired formation and/or stabilization of cilia is a consequence of *zcrb1* loss of function, potentially dysregulating Wnt signaling as a response, in the zebrafish.

**Figure 5:**
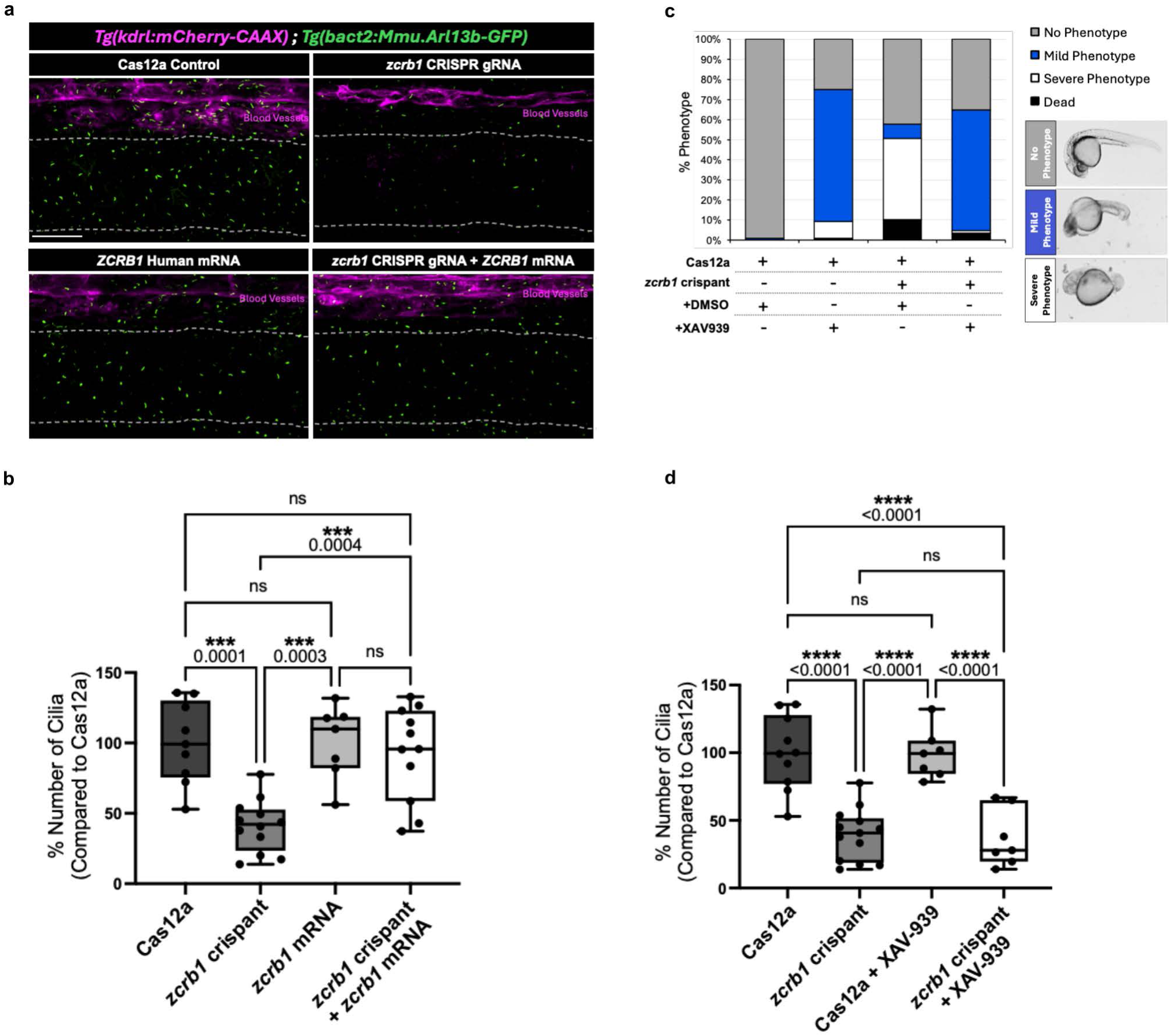
WNT pathway upregulation and cilia downregulation by ZCRB1 loss are conserved in zebrafish a. Cilia formation in Tg(actb2:Mmu.Arl13b-GFP)hsc5Tg transgenic zebrafish as shown in kidney tissue. Cilia are shown in green. b. Quantification of total PC in Cas12a and *zcrb1* crispant embryos, WT embryos injected with *ZCRB1* mRNA, and *zcrb1* crispant embryos injected with ZCRB1 mRNA. Numbers above brackets represent the p-value as measured by an ordinary one-way ANOVA with Tukey’s multiple comparisons test (omnibus p-value=****<0.0001). c. Analysis of gastrulation and body-axis phenotypes in zcrb1 crispants in the presence of the Wnt inhibitor XAV-939 d. Quantification of total PC in zcrb1 crispants in the presence of Wnt inhibitor XAV-939. Numbers above brackets represent the p-value as measured by an ordinary one-way ANOVA with Tukey’s multiple comparisons test (omnibus p-value=****<0.0001).

To determine if the cilia-Wnt signaling axis underlies the gastrulation and body axis phenotypes noted in the *zcrb1* zebrafish Crispants, we treated Cas12a and *zcrb1* Crispant embryos with the Wnt inhibitor, XAV939, and compared phenotypes to DMSO vehicle control-treated embryos^53^. Addition of XAV939 at 20 uM to Cas12a injected embryos led to a mild body axis phenotype, consistent with the known effects of Wnt signaling suppression during development. As noted above (Fig. 4), the *zcrb1* Crispants have marked body axis defects. When treated with XAV939 at 20 uM, the *zcrb1* Crispant injected embryos showed a decrease in the severity of these defects and are phenotypically indistinguishable from the Cas12a injected XAV939 treated embryos (Fig. 5c).

To place Wnt signaling upstream or downstream of cilia, we carried out the same experiment as outlined above, except all treatments were performed on the *Tg(actb2:Mmu.Arl13b-GFP)^hsc5Tg^* transgenic zebrafish background for imaging of cilia (Fig. 5d)^52^ . While we show that treatment of zcrb1 Crispant embryos with the Wnt inhibitor, XAV939, can rescue body axis phenotypes back to the levels of Cas12a embryos treated with XAV939, there is not a corresponding rescue in cilia numbers. Together these data suggest that *zcrb1* splicing of genes regulating cilia formation and maintenance is likely direct, with effects on Wnt signaling secondary to loss of cilia function.

## Discussion

Despite significant strides since the identification of the U11/U12 snRNP associated-proteins over 20 years ago, their biological significance, including *ZCRB1*, and their specific interactions within the minor spliceosome are not fully clear^54,55^. This work provides a focused investigation into the gene regulatory function and significance of ZCRB1, expanding our knowledge of minor spliceosome specific proteins in human biology and disease. We performed CRISPR/Cas9-mediated gene editing on Hek 293 Flp-In cells, creating precise, stable monoallelic knockouts of *ZCRB1*. These knockouts exhibited varying mutant allelic frequencies between 32 and 68%, with corresponding mRNA expression levels measured using quantitative PCR ranging between 44 to 71% of WT values (Table 1, Fig. 1c). Using quantitative PCR (qPCR) and Western blot analyses, we validated the observed differences in mutant allelic frequency and gene expression, allowing exploration of clonal heterogeneity and functional characterization of our mutants. This enabled us to draw comparisons to the natural and non-mutated state as the highly conserved *ZCRB1* has been identified as an essential gene^37,56,57^.

Limited research has shown the highly conserved ZCRB1 to be an RNA-binding protein (RBP) involved in SARS-CoV RNA replication and regulation of cell proliferation in Glioblastoma multiforme (GBM); however, very few studies have examined its role in the minor spliceosome^58^. Recent research by Li et al. functionally characterized bktRNA1 (backward K-turn RNA 1) as a regulator of minor intron splicing through its positive regulation of A8 2′-O-methylation in U12 snRNA^35^. The study demonstrated that btkRNA1 was necessary for the recruitment of ZCRB1 to the U12 snRNA and that a reduction in this interaction caused by depletion of bktRNA1 led to minor intron retention. The study also showed a reduction in minor intron removal efficiency through shRNA-mediated knockdown of *ZCRB1*^35^. In our investigation, we employed differential splicing analysis between human heterozygous *ZCRB1* knockout cells and WT Hek 293 Flp-In cells using rMATs^48^. Employing the same statistical tool used by Li et al study, we saw a dysregulation of both minor and major intron splicing^35^. In our pairwise comparison of cells with heterozygous expression of ZCRB1 and WT cells, we observed a lesser degree of minor intron retention than previously found by Li et al., the latter identified splicing defects in shRNA-treated cells with more than 97% knockdown of ZCRB1 expression (Supplemental Fig. 3a, Supplemental Table 3a)^35^. Our results, however, did show several statistically significant intron retention events, alternate 3’ and 5’ usage, exon skipping events, and mutually exclusive exon events of both MIGs and non-MIGs (Supplemental Fig. 3a, Supplemental Table 3a.). Combined with our results showing significant downregulation of U12 and U1 snRNA, along with several splicing factors necessary for U11/U12 di-SRNP and U1 and U2 snRNP assembly, we can conclude that ZCRB1 has effects on global splicing (Fig. 2b., Supplemental Table 2c).

Beyond splicing, RNA-seq permitted a comprehensive analysis of the effect of heterozygous loss of ZCRB1 on gene expression. Despite the incomplete loss of expression in zebrafish, we saw the significant differential expression of 4582 genes involved in a multitude of essential cellular processes (by gene ontology analyses). Notably the significantly downregulated biological processes included, ribonucleoprotein complex biogenesis and RNA processing. This finding underscores not only the impact of *ZCRB1* loss on splicing but also highlights its broader role in various aspects of RNA metabolism, including the regulation of other major and minor spliceosome-associated splicing factors, snRNPs, and related components. Gene ontology analysis revealed an upregulation in signaling and regulatory processes related to growth, differentiation, and development (Extended Figure 2, Supplemental Table 2b). This, combined with increased signaling pathways for cell communication and signal transduction, prompted us to explore the impact of *ZCRB1* knockdown on Wnt signaling -a crucial pathway for proliferation, differentiation, and tissue development^59^. Employing both our heterozygous *ZCRB1* knockout Hek293 Flp-In cells and the zebrafish *zcrb1* Crispant model, we saw an upregulation of Wnt ligands and a downregulation of Wnt inhibitors (Figure 2c, Supplemental Table 2c, Figure 4f, Supplemental Table 4c) Notably, an increased expression of the Wnt ligands Wnt11 and Wnt9 in both organisms was observed, indicating Wnt pathway overactivation associated with heterozygous loss of *ZCRB1* (Figure 2c, Supplemental Table 2c, Figure 4f, Supplemental Table 4c). Additionally, there were transcriptional differences in the Wnt signaling pathways in zebrafish, also manifested phenotypically through disrupted gastrulation and altered dorsal-ventral body axis polarity. It is established that Wnt signaling in zebrafish activates the Spemann organizer, which plays a crucial role in dorsal-ventral patterning by inhibiting BMP (Bone Morphogenetic Protein) signaling^60,61^. This interaction helps establish a gradient of Wnt signaling activity, which is essential for proper embryonic patterning^60,61^. Overexpression of Wnt ligands causes posteriorization in zebrafish embryos, leading to anterior defects such as the absence of eyes and forebrain, a phenotype observed in our *zcrb1* mutants^60^. Moreover, overactivation of Wnt signaling during embryogenesis also negatively affects gastrulation^62^. These observations suggest a direct link between the transcriptional effects of the Wnt overexpression and its impact on zebrafish embryonic development. The ability of WT *ZCRB1* to rescue these gastrulation and body axis phenotypes suggests a role of the downstream functional role of z*crb1* in Wnt signaling.

Despite seeing an upregulation of Wnt signaling in both our human and zebrafish RNA-seq data, as well as in the observed zebrafish phenotypes, there remains a notable gap in the scientific literature connecting dysregulation of minor spliceosome specific factors and alterations in Wnt signaling. Upon revisiting the RNA-seq data with a focus on MIG expression, we found that 28% of these genes were significantly differentially expressed. Among these, 73% were downregulated (Figure 3a, Supplemental Figure 3a). Importantly, none of the differentially expressed MIGs were canonical Wnt genes. Our focused gene ontology analysis showed an enrichment of differentially expressed MIGs involved in processes most notably associated with primary cilia assembly, maintenance, and function (Figure 3b). Primary cilia are vital for cellular signaling, and their proper formation is crucial for maintaining cellular functions. The observed significant downregulation of MIGs involved in essential ciliary processes including but not limited to IFT, the formation of the BBsome, and the formation of the MKS and NPHP gene complexes, suggests that ZCRB1 plays a critical role in the splicing and expression of these genes and thus is necessary for ciliogenesis and function. Moreover, heterozygous loss of *ZCRB1* may have disease implications as many of the top downregulated genes in the heterozygous *ZCRB1* knockout cells, such as *CEP290, TMEM67, RPGRIP1L*, and *IFT88*, are causative agents of multiple ciliopathies^63–66^. Additionally, the RPE1 model showed a significant loss of cilia formation coordinated with loss of minor-intron-containing ciliogenesis factors including IFT88, RABl3b, and CEP41 (Figure 3d-f). Loss of Ift88 in RPE1 cells has been shown in previous studies to prevent cilia formation, supporting a role of ZCRB1 in ciliogenesis through modulation of *IFT88* expression^67^. Through CRISPR/Cas12a targeting of transgenic GFP cilia-labeled zebrafish with *zcrb1* gRNA, we were able to show that similarly to the experiments in human RPE-1 cells shown in this work, incomplete loss of *zcrb1,* is sufficient for loss of primary cilia formation in zebrafish. The rescue of PC formation in WT *ZCRB1* mRNA further signifies this functional role for *zcrb1*. Dysregulation of primary cilia has been seen in human cells in both the loss of the minor spliceosome-specific *SCNM1* gene and in zebrafish with minor spliceosome-specific U4atac snRNA, further supporting a role of *ZCRB1* in primary ciliogenesis^28,29^. These findings align with the known functions of ZCRB1 in the minor spliceosome and extend its role to ciliary biology, providing new insights into the molecular mechanisms underlying ciliary dysfunction.

A key finding from this work is that treatment with a Wnt inhibitor, XAV939, resulted in the recovery of gastrulation and dorsal-ventral patterning defects in our zebrafish mutants. However, this treatment did not restore cilia formation, indicating that the observed effects of Wnt signaling are likely a consequence of ciliary loss rather than a direct cause. There are several studies that show that Wnt and cilia have a complex feedback mechanism in which they regulate each other, while others suggest that Wnt signaling is not required for ciliogenesis^50,68^. Our current model suggests that heterozygous loss of ZCRB1 in vertebrate cells causes the mis-splicing of ciliary genes, causing disrupted Wnt signaling and further downstream developmental abnormalities (Figure 6). While this work does not fully resolve the controversy surrounding the relationship between Wnt signaling and ciliogenesis, it provides valuable insights that may clarify aspects of this complex interplay while providing context for the function of the minor spliceosome in these processes.

**Figure 6:**
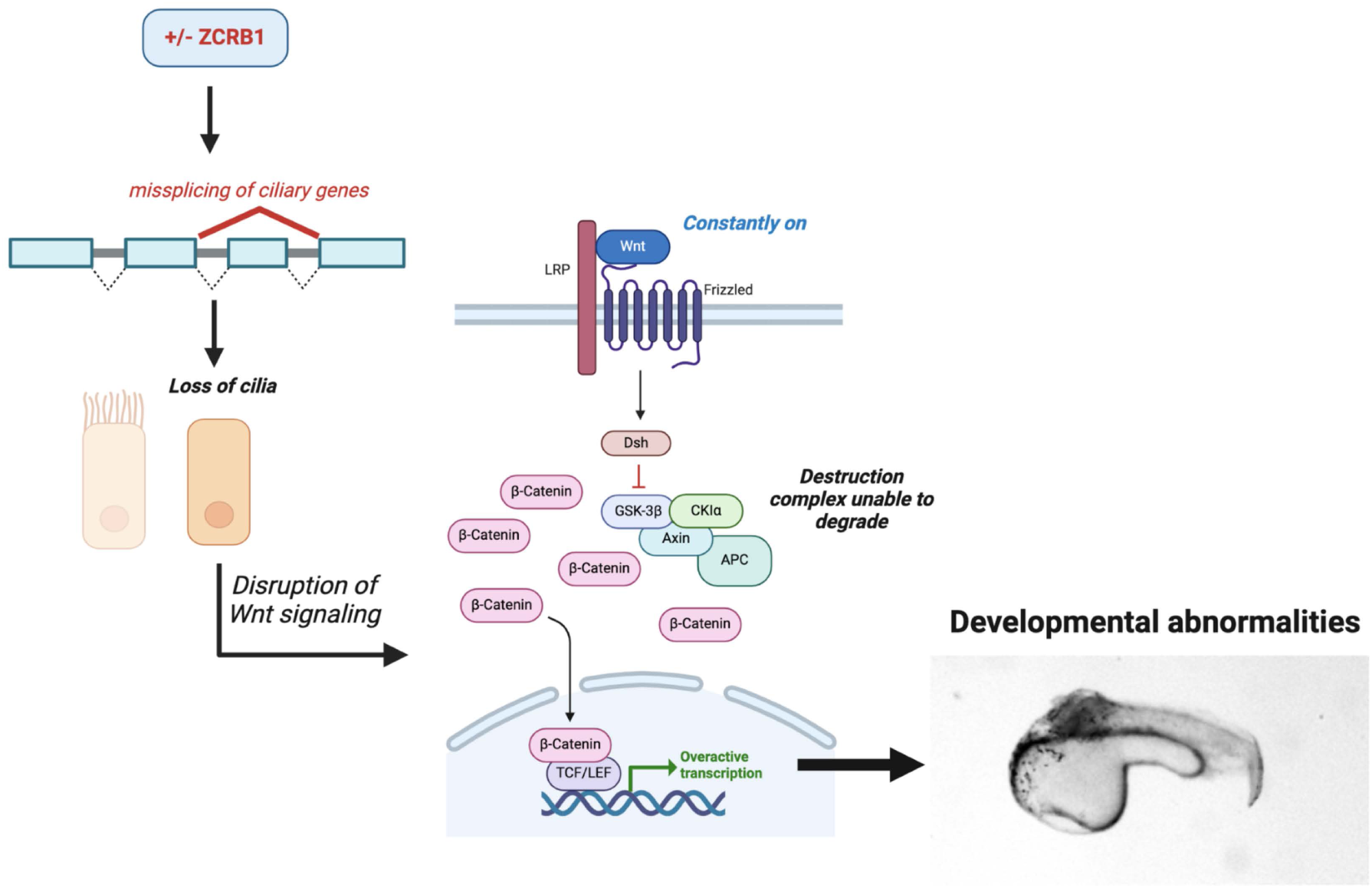
Model depicting role of partial ZCRB1 on ciliogenesis, Wnt signaling, and vertebrate development.

Taken together this work provides the first detailed functional characterization of ZCRB1 in splicing and beyond using the first cellular and zebrafish model of *zcrb1* partial and complete loss, offering new insights into its role and biological significance while highlighting complex mechanistic interactions between ciliogenesis and Wnt signaling, and expanding our knowledge of the biological significance of the minor spliceosome.

## METHODS

### *Homo sapiens* cell culture experiments

Hek293 Flp-In T-REx cells were cultured in Dulbecco’s modified Eagle’s medium (DMEM) (Gibco) and supplemented with 10% FetalGro bovine growth serum (BGS), 5% minimum essential medium nonessential amino acids (100×, Gibco), 5% penicillin, streptomycin (Gibco), and L-glutamine (Gibco). Additionally, 5 μg mL−1 of blasticidin and 100 μg mL−1 of Zeocin were added to the medium for cell line maintenance. Cells were maintained at 37°C with 5% CO_2_. For CRISPR-Cas9 editing cells were grown to 70-80% confluency. Predesigned Alt-R^TM^ CRISPR-Cas9 single guide RNAs (sgRNAs), RNA oligonucleotide containing both crRNA and tracrRNA regions, were selected for the ZCRB1 gene from Integrated DNA Technologies (IDT). Two sgRNAs were selected based on high predicted editing performance and lower off-target risk. The Design ID, strand (+ or -), and sequence for sgRNA 1 and 2 are as written, respectively: HS.Cas9.ZCRB1.1.AA, +, GTACAAGTCATTGTTTGTCA; HS.Cas9.ZCRB1.1.AB-, GCAAGCATTGCTATTGACAA. Hek 293 Flp-In T-REx cells were electroporated with a 1.2.to 1 molar ratio of sgRNA to CRISPR/Cas9 (GeneArt™ Platinum™ Cas9 Nuclease, Thermofisher Scientific) using the Neon™ Transfection System (ThermoFisher Scientific). We expanded individual CRISPR-edited cells through a process of clonal expansion to ensure monoclonality. Following expansion, edited cells were assessed for ZCRB1 expression at the mRNA and protein levels, by qPCR and Western blot analysis, respectively. Whole Genome Sequencing (WGS) analysis facilitated the identification of on and off-target events.

### Whole Genome Sequencing and Analysis

Libraries were constructed from 600ng of DNA using the KAPA Hyper PCR-free Library Prep Kit (KAPA Biosystems) on a SciClone NGS instrument (Perkin Elmer). Genomic DNA was fragmented using a Covaris focused-ultrasonicator with a target insert size of 350 base pairs followed by a paramagnetic bead cleanup for size selection. The library concentrations were determined using qPCR (Kapa Biosystems) and pooled. Pooled libraries were sequenced to generate paired end reads of 150 bases using the S4 300 Cycle kit with XP workflow on the Illumina NovaSeq 6000. Illumina’s bcl2fastq2 software was used for base calling and demultiplexing to create sample specific FASTQ files.

Variant Calling. The samples were analyzed on a DRAGEN BioIT processor running software version 4.0.3. Sequences reads were aligned to the GRCh38 reference genome witAlignments were generated in CRAM format. Structural variants were called along with small and copy number variants. BCFtools was used for the manipulation and parsing of CRAM files^69^. Functional characterization of single nucleotide polymorphisms was performed using SNPeff^70^.

### Protein extraction SDS PAGE and Immunoblotting blotting (IB)

Hek293 Flp-In T-REx cells were lysed in adequate amount 1X RIPA buffer (9806, cell signaling (CS)) supplemented with 200mM PMSF and 1X protease/phosphatase inhibitor (5872, CS) and passed through 1ml syringes thrice (30G) for efficient lysis. Supernatants were collected after performing the centrifugation at 4°C for 20 minutes at 20,000G. Protein estimation was performed by Bradford assay (5000006, Bio-Rad (BR)) and adequate amount was loaded on SDS PAGE gradient gels (3450123, BR). The samples were dissolved in sample loading buffer (1610791, BR) supplemented with reducing agent (1610792, BR) and subjected to run at constant 150V in presence of XT MES running buffer (1610789, BR). Separated proteins were blotted on a polyvinylidene fluoride (PVDF) membrane by semi-dry transfer at 25V for 30 minutes. The membranes were blocked with 5% milk in PBST (0.1% triton in PBS) for 30 minutes and incubated with primary antibodies either for 2 hr at room temperature (RT) or overnight in the cold room at constant shaking. Secondary HRP conjugated antibodies were incubated for 1 hr at RT, following the antibody incubation membranes were washed thrice with PBST for 5 minutes and proteins levels were detected by chemiluminescence (34577, BR). The following primary antibodies were used in this study ZCRB1 (25629-1-AP, proteintech), LRP6 (2560, CS), Phospho-LRP6 Ser1490 (2568, CS), IFT88 (13967-1-AP, proteintech), GSK3ß (9315, CS), ß-actin (4967, CS), RABL2B (11588-1-AP, proteintech), and CEP41 (PA5-103727, Invitrogen). HRP conjugated anti-Rabbit IgG (7074, CS) was used as secondary antibody in this study.

### RNA isolation, cDNA synthesis, and quantitative real time PCR (qPCR)

RNA was extracted using RNeasy mini kit (74104, Qiagen) by following the manual instructions. The extracted RNA was treated with TURBO DNAase (AM1907, Thermo Fisher Scientific) followed by cDNA synthesis using iScript advanced cDNA synthesis kit (1725037, BR). The qPCR was performed by using iQ SYBR green supermix (1708880, BR) by following the manual instructions in a c1000 thermocycler (CFX96, BR). The qPCR data was normalized to a house keeping gene called GOGLA5. For each reaction three technical replicates and for each experiment atleast four biological replicates were performed. The following primers were used for amplification *ZCRB1* (NM_033114) forward primer (FP): CTCCAAGTAAGAGCACAGTGT *ZCRB1* reverse primer (RP): CAACCCCTTTACTCTTCCTGG *GOLGA5*(NM_005113.4) FP: CGAACAGCAGATGAACTCCG, *GOLGA5* RP: AGCTTTGCGAACTTTTCCGT.

### Small interfering RNA (siRNA) transfections, Immunofluorescence, and microscopy

Transient depletion of *ZCRB1* in RPE1 cells was performed by electroporating an equimolar cocktail of siRNA 2 and 3 (150pM). The siRNA treatment was done for 100hr by electroporating the cells twice at 0hr and at 50hr. The electroporation was performed by neon electroporation system (MPK5000, Invitrogen). The *ZCRB1* siRNAs were obtained from IDT targeting the CDS region of *ZCRB1* mRNA with the following sequence, siRNA2: 5’ -GCAAGCAUUGCUAUUGACAAUGGAA-3’, and siRNA3: 5’-CCCUCAACAUCAGAUGAUUCAAGAC-3’. To induce the formation of PC the cells were serum starved (0.5%) for 48 hr (Kengo Takahashi et al., 2018) before performing the IF and IB. TYE 563 transfection control, HPRT-s1 Positive control and Negative control siRNAs from the provider were used to validate the efficiency and specificity of siRNA setup. For IF the cells were seeded to achieve 80% confluency prior to fixation on glass coverslip (12541007, Fisher Scientific). After ice-cold methanol fixation for 20min at – 20°C, cells were rinsed thrice with 1X PBS and permeabilized with 0.1% triton X-100 in PBS (PBS-T) for 20min at RT. Followed by blocking with 3% BSA (A2153, Sigma) in PBS-T at RT. The samples were incubated with CEP120 (generated in Moe R Mahjoub Lab) and Acetylated tubulin (T6793, Sigma) primary antibodies overnight in PBS-T. After 3 PBST washes the samples were incubated with alexa fluor dye conjugated secondary antibodies (A-11008, A-11005, Thermo) for 1 hr at RT. The nuclei were stained with DAPI (62248, Thermo) or Hoechst (H3570, Life technologies) and the samples were mounted with cytoseal XYL (8312-4, epredia). Images were captured with Zeiss LSM 880 confocal microscope and processed with Fiji-imageJ software.

### CRISPR/Cas12a and mRNA injections into zebrafish Embryos

Guide RNAs (gRNAs) to zebrafish *zcrb1* were identified using CHOPCHOP. gRNA 5’ –TTCC GAC AGT TAT GCT CAA AAT ATG GCA –3’

10 nmol of Alt-R A.s. Cas12a crRNA was ordered from IDT (as a pre-synthesized gRNA) for injection with 100ng/pl Cas12a (Cpf1) for editing (IDT Alt-R A.s. Cas12a. (Cpf1) Ultra, #10001272). CRISPR/Cas12a mixtures were injected into wild type AB* zebrafish embryos at the 1 cell stage ^51,71^.

Human *ZCRB1* mRNA was generated by in vitro transcription of pcDNA-DEST40 reporter c^3^onstructs containing the coding transcript of ZCRB1 using the mMESSAGE mMACHINE™ T7 Transcription Kit. mRNA was treated with TURBO DNAase (AM1907, Thermo Fisher Scientific) and purified by ethanol precipitation with 3M sodium acetate before injection into wild type AB* or co-injected with gRNA into wild type AB* zebrafish embryos at the 1 cell stage^72^.

Mixtures to be injected as 1 nl boluses into zebrafish 1 cell stage embryos included:

#### *zcrb1* gRNA

0.85 µL of 10 nmol individual stock gRNA

1.0 µL of 100 ug stock Cas12a (Cpf1)

1.95 µL 2M KCl

5.0 µL H2O

1.0 µL Phenol red

#### *zcrb1* gRNA *+* Wild-Type (WT) *ZCRB1* mRNA

0.85 µL of 10 nmol individual stock gRNA

1.0 µL of 100 ug stock Cas12a (Cpf1)

1.95 µL 2M KCl

0.25 µL H2O

5.0 µL WT *ZCRB1* mRNA

1.0 µL Phenol red

#### Cas12a (Cpf1) only control

1.0 µL of 100 ug stock Cas12a (Cpf1)

1.95 µL 2M KCl

6.05 µL H2O

1.0 µL Phenol red

#### WT *ZCRB1* mRNA only

1.95 µL 2M KCl

2.05 µL H2O

5.0 µL WT *ZCRB1* mRNA

1.0 µL Phenol red

WT *ZCRB1* mRNA and phenol red were added following a 10-minute incubation of the gRNA/Cas12a mixture at 37°C. The mixture was then kept on ice until microinjection into the zebrafish embryos at the one cell stage.

### Imaging of cilia and *wnt* transgenic zebrafish reporters

Transgenic zebrafish lines used in this work: *Tg(OTM:d2EGFP)^kyu1Tg^*; *Tg(actb2:Mmu.Arl13b-GFP)^hsc5Tg^*^52,73^.

*Tg(actb2:Mmu.Arl13b-GFP)^hsc5Tg^* or *Tg(OTM:d2EGFP)^kyu1Tg^* adult zebrafish were crossed and injected with the gRNA/Cas12a mixtures described above. Following injection, the embryos were housed at 28°C for 28 hours for the assessment of phenotypes. In all cases, the embryos were allowed to mature for ∼3 hours post injection and unfertilized embryos removed to aid in accurately assessing gastrulation and live/death ratios following injections. Embryos were grown for approximately 28 hours prior to phenotypic and genotypic analysis.

Imaging was done using a Ti2 Nikon microscope with CSU-W1 confocal scanner, 20x or 40x APO-Plan objective, and a Fusion camera to image cilia and Wnt activation within zebrafish embryos. A Z-stack step size of 0.4 uM was used for all acquisitions. At 24 hpf, embryos were dechorionated using Dumont Tweezers, Style 55 (Electron Microscopy Sciences; #72707-01). The embryos were immobilized using MS-222 and embedded in 1% UltraPure low melting point agarose (ThermoFisher #16520050) and fish water for live imaging.

### Genotyping and Fragment Analysis of *zcrb1* gRNA injected embryos

Embryos were placed individually in PCR tubes and genomic DNA (gDNA) extracted using equal volumes of extraction solution (Sigma Aldrich; #SLCQ0691) and tissue preparation solution (Sigma Aldrich; #SLCH8297). Samples were incubated in a SimpliAmp Thermal Cycler (Thermo Fisher Scientific) for 15 min at 55°C, followed by 15 min at 95°C to allow for tissue digestion. Primers flanking the *zcrb1* gRNA sites were used to generate small amplicons for analysis of CRISPR cutting efficiency using an Agilent 5300 Fragment Analyzer^51^. Primers:

gRNA #2: Fw-5՝-AAGGATCATGTTTTGCATGTTG

Rv-5՝-ATGATCATCCACTTGGCAGTTA

gRNA #3: Fw-5՝-GGCCTAAAAATAACAAAGCATCA

Rv-5՝-TCAAGCCATGCATTGACATAAA

gRNA #4: Fw-5՝-GATGAGTAAAGGAGTGGCGTTC

Rv-5՝-CACACCTGAACCAGCTAATCAA

### RNA sequencing and Analysis

RNA was extracted using the RNeasy system (Qiagen) for Human Cell lines and for Zebrafish RNA concentration was measured by NanoDrop (ThermoFisher Scientific). Total RNA integrity was determined using Agilent Bioanalyzer or 4200 Tapestation. Library preparation was performed with 1ug of total RNA. Ribosomal RNA was removed by an RNase-H method using RiboErase kits (Kapa Biosystems) for Human cell lines and by a by a hybridization method using Ribo-ZERO kits (Illumina-EpiCentre for Zebrafish samples. mRNA was then fragmented in reverse transcriptase buffer and heating to 94 degrees for 8 minutes. mRNA was reverse transcribed to yield cDNA using SuperScript III RT enzyme (Life Technologies, per manufacturer’s instructions) and random hexamers. A second strand reaction was performed to yield ds-cDNA. cDNA was blunt ended, had an A base added to the 3’ ends, and then had Illumina sequencing adapters ligated to the ends. Ligated fragments were then amplified for 12-15 cycles using primers incorporating unique dual index tags. Fragments were sequenced on an Illumina NovaSeq-6000 using paired end reads extending 150 bases. Basecalls and demultiplexing were performed with Illumina’s bcl2fastq software with a maximum of one mismatch in the indexing read. RNA-seq reads were then aligned to the Ensembl release 101 primary assembly with STAR version 2.7.9a1. Gene counts were derived from the number of uniquely aligned unambiguous reads by Subread:featureCount version 2.0.32. Isoform expression of known Ensembl transcripts were quantified with Salmon version 1.5.23. The ribosomal fraction, known junction saturation, and read distribution over known gene models were quantified with RSeQC version 4.04. TMM normalization size factors were calculated using the R/Bioconductor package EdgeR^74^. Ribosomal genes and genes not expressed in the smallest group size minus one samples greater than one count-per-million were excluded from further analysis. The TMM size factors and the matrix of counts were then imported into the R/Bioconductor package Limma^75^. Differential expression analysis was performed using Limma to analyze for differences between controls and CRISPR edited Human and Zebrafish samples, respectively. Gene set enrichment was performed using GAGE^40^. Statistically significant genes were considered to have an adjusted p-value (FDR) <0.05 and a log2FC <-.58 or > .58. GO terms were filtered for pathways scored with adjusted p-values (FDR) <0.05.

### Differential Splicing Analysis

Alternative splicing analysis for both Human and Zebrafish samples was performed using the rMATS (turbo0.1) on a cloud computing platform. BAM files generated from sequencing alignments as previously described were used for analysis. Alternative splicing events, including exon skipping (SE), mutually exclusive exons (MXE), retained introns (RI), alternative 5’ splice sites (A5SS), and alternative 3’ splice sites (A3SS) were identified for each sample and then compared across conditions to detect differences in alternative splicing events. Alternative splicing events were filtered for events containing adjusted p-value (FDR) < 0.05 and delta PSI > 0.05. Gene annotation for major and minor Human intron-containing genes was performed using list of human introns as described by Norppa et al^18^ . Functional enrichment of MIGs was performed using ShinyGO v0.80^77^. GO terms were filtered for pathways scored with adjusted p-values (FDR) <0.05.

### Statistical Analysis

Statistical analyses for figure panels 1c, 3f, 5b and 5d were performed using GraphPad Prism 10. Data normality was determined by using the D՝Agostino-Pearson omnibus test. Statistical analyses, post-hoc tests and P-values are all described in corresponding figures and figure legends. Significance was determined by a P-value of 0.05 or less. The p-values associated with the RNA sequencing data were calculated with the Limma software. Fischer’s exact testing was performed using the scipy.stats module from the SciPy library^78^.

## Acknowledgments

We are thankful to all the members of Pavlovic Djuranovic, Stratman and Djuranovic labs for their help on our study and writing of the manuscript. We are thankful to Dr. Mikko Frilander, Dr Juan Alfonzo and Dr. Susan Dutcher for their valuable comments and careful reading of the manuscript. The work in Djuranovic Lab was sponsored by NIGMS R01GM136823 and R01GM112824, Chen-Zuckerberg Neurodegeneration Initiative and Siteman Cancer Center Investment Award. Work in Stratman Lab was sponsored by NIGMS R35GM137976. We thank the Genome Technology Access Center at the McDonnell Genome Institute at Washington University School of Medicine for help with genomic analysis. The Center is partially supported by NCI Cancer Center Support Grant #P30 CA91842 to the Siteman Cancer Center from the National Center for Research Resources (NCRR), a component of the National Institutes of Health (NIH), and NIH Roadmap for Medical Research. This publication is solely the responsibility of the authors and does not necessarily represent the official view of NCRR or NIH.

---

**Extended Data Fig. 1:**
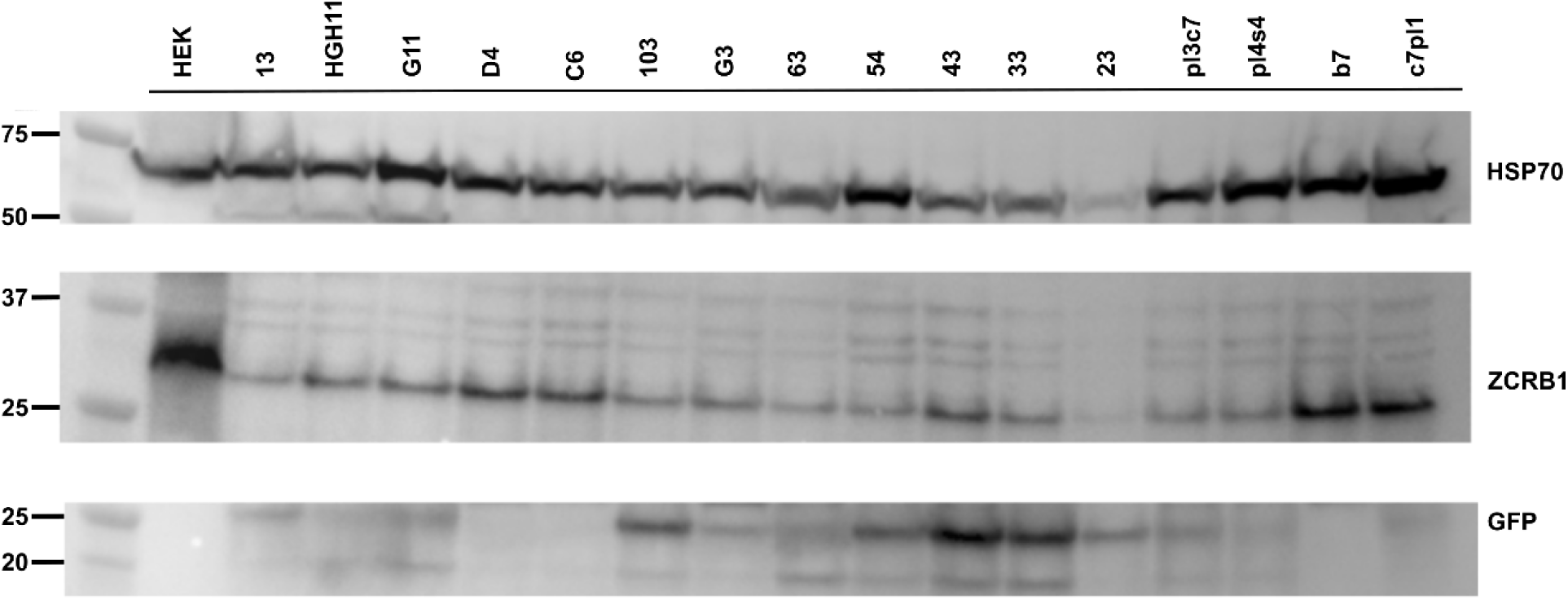
Generation of heterozygous *ZCRB1* knockout cell lines. Representative Western blot image showing ZCRB1 protein expression in individually CRISPR/Cas9 edited cell lines and in the parental HEK293 Flp-In cell line. GFP expression is shown a positive marker of genomic editing and HSP70 is represented as a loading control.

**Extended Data Fig. 2:**
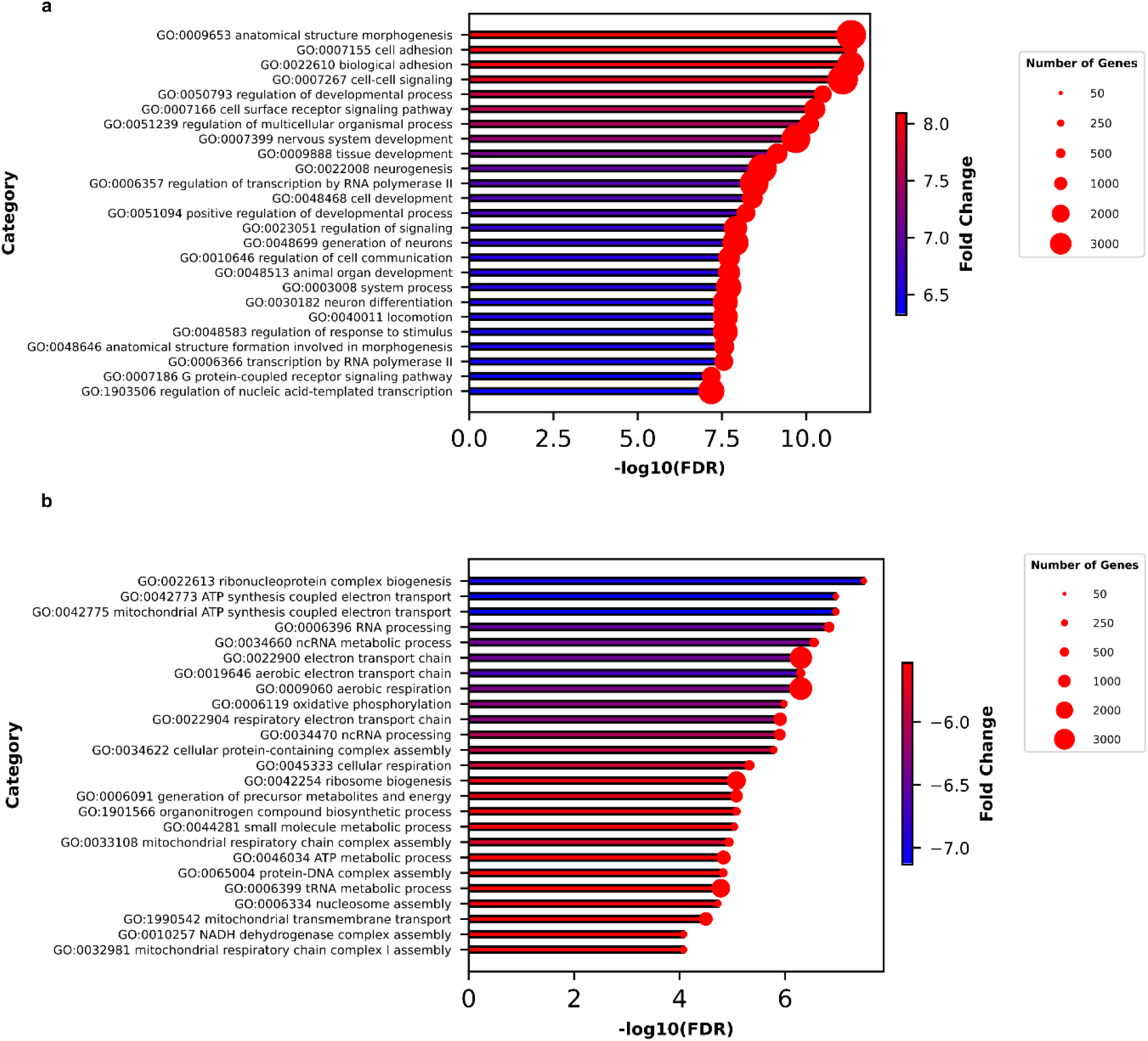
FDR significant enriched biological processes (BP) identified in heterozygous *ZCRB1* knockout cell lines versus WT. a. Bar-plot representing the top 25 upregulated BP. Gene ontology (GO) terms and IDs are located on left hand side of bars. The length of bars is proportional to the FDR (adjusted p value) and the shading dictates the degree of linear fold change shown by the color bar. The diameter of the red dots on the end of the bars represents the number of genes present in each identified enriched pathway. b. Bar-plot representing the top 25 downregulated BP. Figure notations are as previously described.

**Extended Data Fig. 3:**
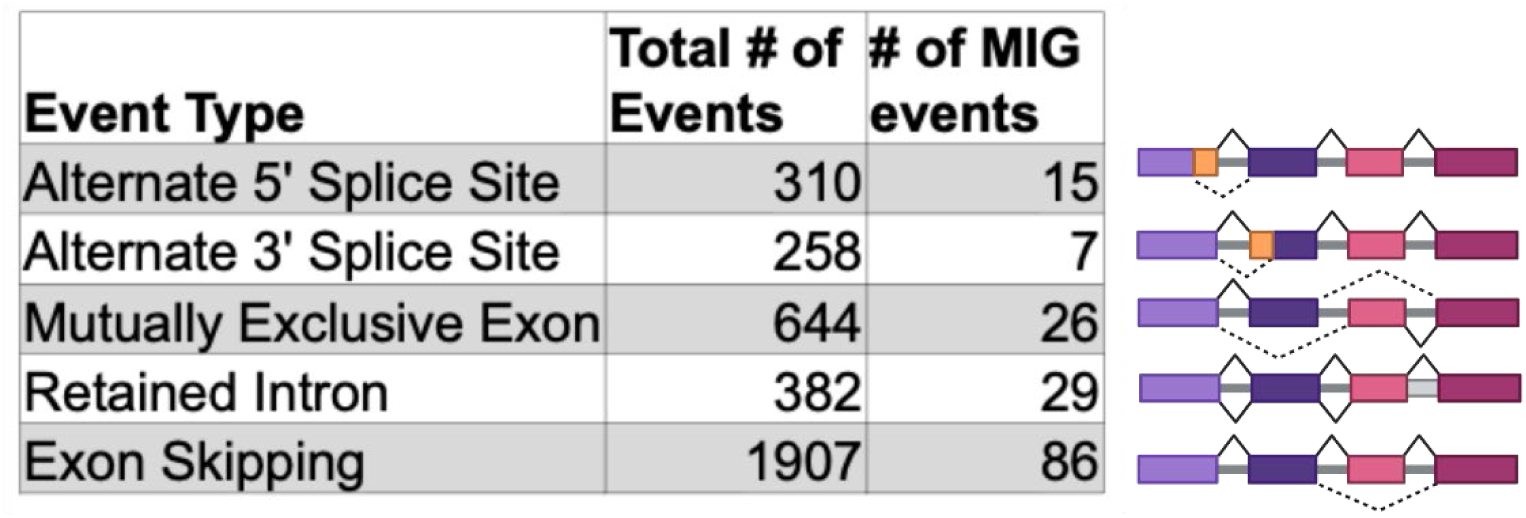
FDR significant alternative splicing events categorized by MIG and non-MIG. Schematics are representative of the labeled events. Colored bars represent exons, straight lines represent introns, and bent lines represent possible splice isoforms.

**Supplemental Fig. 1:**
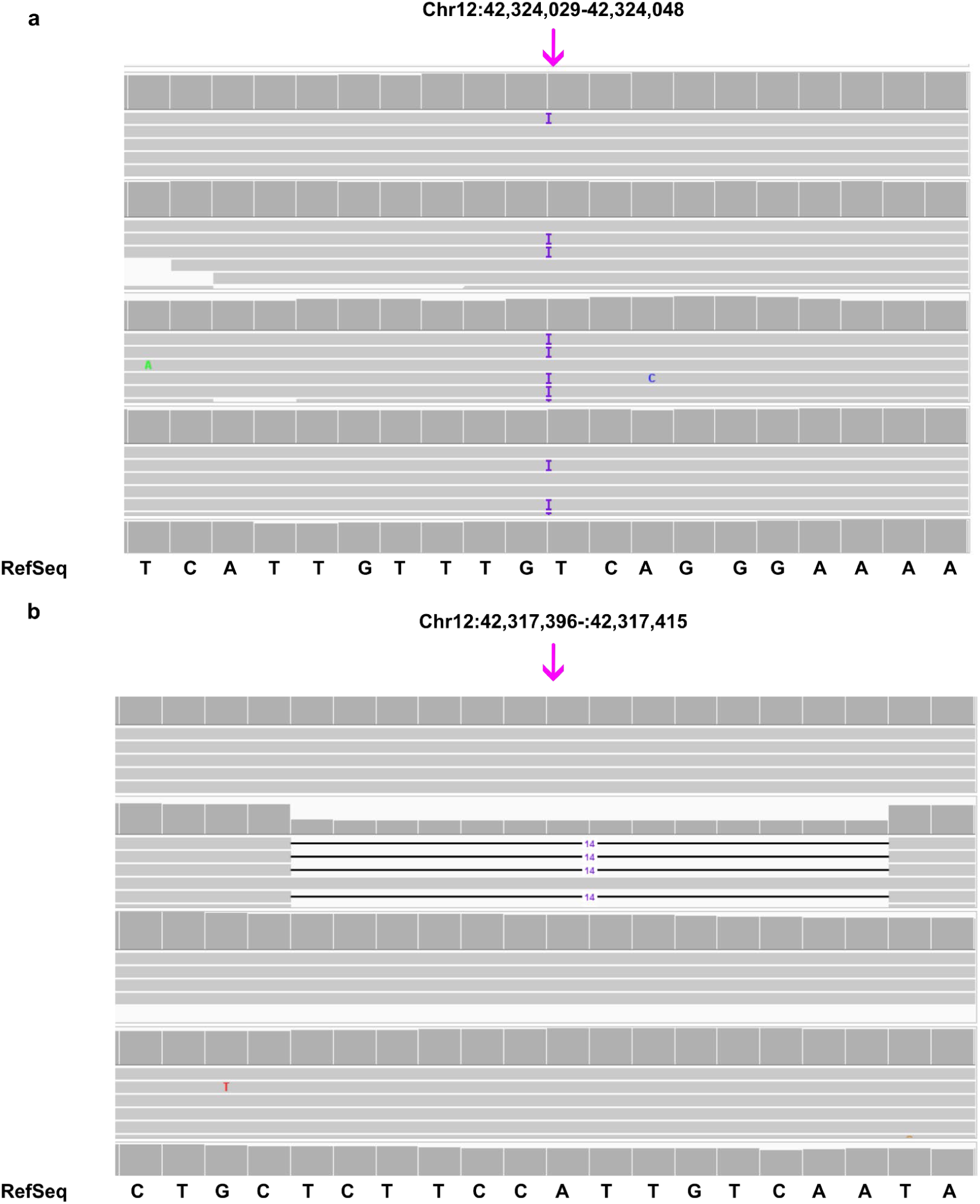
CRISPR/Cas9 mediated editing of *ZCRB1* a. IGV snapshot showing the genomic region targeted by sgRNA 1, highlighting a 1 bp insertion mutation. The location of the mutation is indicated by an arrow. The reference sequence is indicated by bold text. b. IGV snapshot showing the genomic region targeted by sgRNA 2, highlighting a 14 bp deletion mutation. The location of the mutation is indicated by an arrow. The reference sequence is indicated by bold text.

**Supplemental Fig 2:**
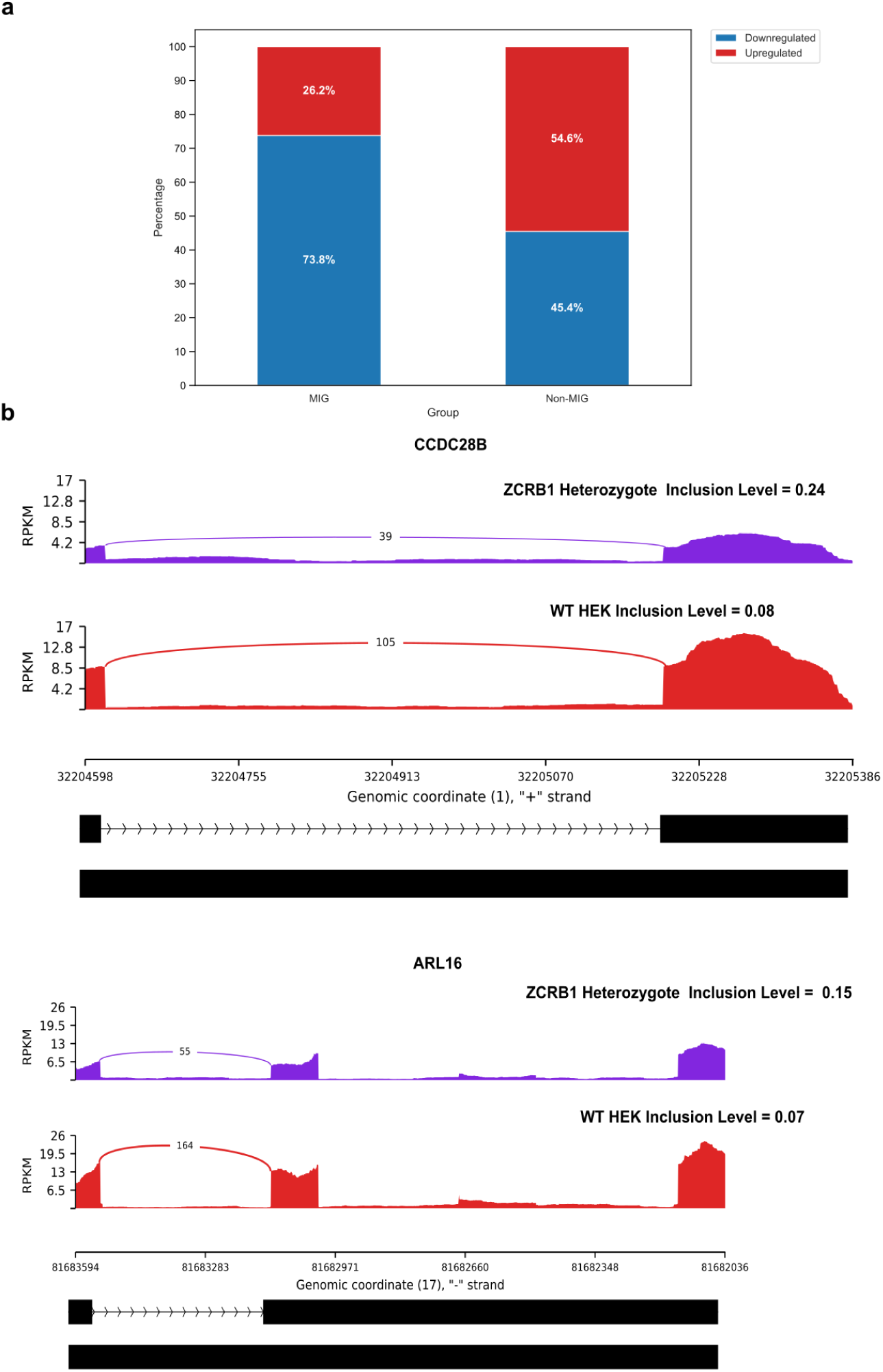
Mis-regulation and splicing of MIG expression in heterozygous ZCRB1 knockout cells. a. Barplot representing the percentage of DE MIG and non-MIG genes as identified by RNA-seq analysis. b. Sashimi plots representing the level of intron retention in MIG genes *ARL16* and *CCDC28B*, respectively in ZCRB1 Heterozygotes versus WT cells. The plot shows the distribution of RNA-seq reads across exons and introns, with the y-axis representing RPKM (Reads Per Kilobase of transcript per Million mapped reads). The exon junction counts are represented in the arcs connecting two exons spanning the retained intron.

**Supplemental Figure 3:**
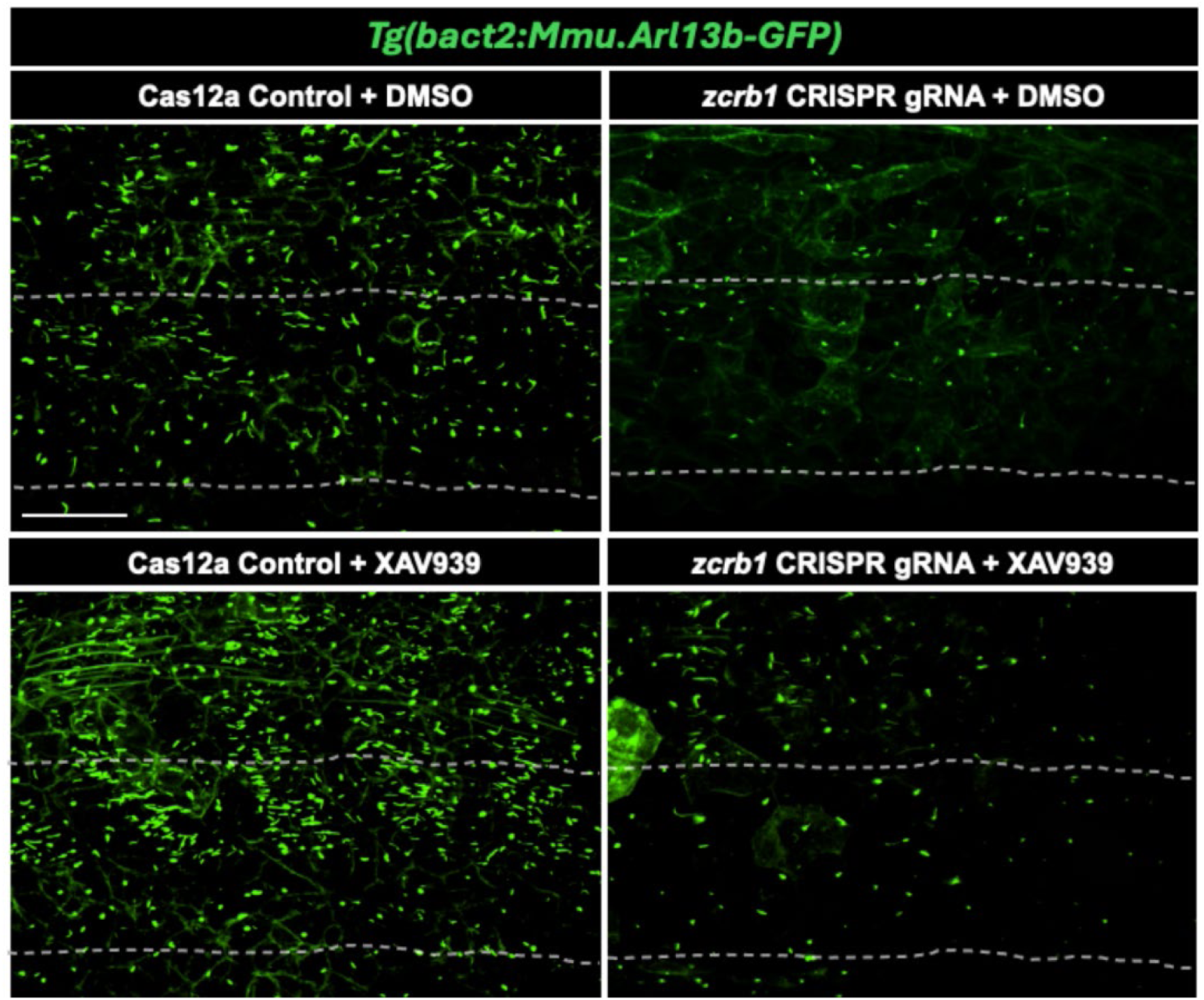
Cilia formation in Tg(actb2:Mmu.Arl13b-GFP)hsc5Tg transgenic zebrafish as shown in kidney tissue. Cilia are shown in green.

## Notes

### Competing Interest Statement

The authors have declared no competing interest.

